# A Computational Toolset for Rapid Identification of SARS-CoV-2, other Viruses, and Microorganisms from Sequencing Data

**DOI:** 10.1101/2020.05.12.092163

**Authors:** Shifu Chen, Changshou He, Yingqiang Li, Zhicheng Li, Charles E Melançon

**Affiliations:** Shenzhen Institutes of Advanced Technology, Chinese Academy of Sciences, Shenzhen, Guangdong, 518055, China; HaploX Biotechnology, Shenzhen, Guangdong, 518057, China

**Keywords:** SARS-CoV-2, viruses, microorganisms, identification, k-mer

## Abstract

In this paper, we present a toolset and related resources for rapid identification of viruses and microorganisms from short-read or long-read sequencing data. We present fastv as an ultra-fast tool to detect microbial sequences present in sequencing data, identify target microorganisms, and visualize coverage of microbial genomes. This tool is based on the k-mer mapping and extension method. K-mer sets are generated by UniqueKMER, another tool provided in this toolset. UniqueKMER can generate complete sets of unique k-mers for each genome within a large set of viral or microbial genomes. For convenience, unique k-mers for microorganisms and common viruses that afflict humans have been generated and are provided with the tools. As a lightweight tool, fastv accepts FASTQ data as input, and directly outputs the results in both HTML and JSON formats. Prior to the k-mer analysis, fastv automatically performs adapter trimming, quality pruning, base correction, and other pre-processing to ensure the accuracy of k-mer analysis. Specifically, fastv provides built-in support for rapid SARS-CoV-2 identification and typing. Experimental results showed that fastv achieved 100% sensitivity and 100% specificity for detecting SARS-CoV-2 from sequencing data; and can distinguish SARS-CoV-2 from SARS, MERS, and other coronaviruses. This toolset is available at: https://github.com/OpenGene/fastv.

## INTRODUCTION

The COVID-19 pandemic has spread to over 200 countries and territories, and has made a terrible impact on lives and economies worldwide [1-3]. Based on the current pandemic situation and many research reports [4], COVID-19 may continue to spread for a long period of time, and may eventually become a flu-like seasonal outbreak [5]. Under these circumstances, it is important to develop new technologies for rapid detection of COVID-19.

Nucleic acid sequencing is a key technology for identifying and studying SARS-CoV-2, the causative agent of COVID-19. Metagenomic and metatranscriptomic next-generation sequencing (mNGS) are powerful tools to study the genetic composition and function of microbial populations, as well as to analyse the relationship between microorganisms and their host or environment [6, 7]. Since the first clinical application of metagenomic sequencing (mNGS) for the diagnosis of leptospirosis in 2014 [8], mNGS has been used widely for the identification and diagnosis of new and rare pathogens.

NGS technology has also played an important role in COVID-19 diagnosis and research. The earliest COVID-19 case, which was initially diagnosed as pneumonia caused by an unknown pathogen, was identified due to the presence of SARS-like sequences found using mNGS [9]. The first complete genome of SARS-CoV-2 (GenBank: MN908947) reported on January 11, 2020 was assembled using NGS data [10]. Soon after, the whole-genome sequence of SARS-CoV-2 was also obtained by mNGS using the Oxford Nanopore platform supplemented with Sanger sequencing [11]. The rapid acquisition and publication of the SARS-CoV-2 genome was essential to design fluorescent PCR probes for COVID-19 nucleic acid detection kits.

With the viral genome in hand, we can now explore the possibility of using mNGS directly as a detection method to determine whether SARS-CoV-2 RNA is present or absent in a sample. In theory, a simple and straightforward approach would be to first map sequencing reads obtained from the sample to the viral genome using common aligners such as BWA [12] or Bowtie2 [13], and then to analyze the alignment results to determine the coverage of the viral genome and the number of properly mapped reads. However, in practice, such an alignment-based method is prone to problems stemming from both false positives and false negatives. On one hand, some viruses have genomes very similar to SARS-CoV-2, which can lead to false positive results. For example, the genome of bat coronavirus RaTG13 is sufficiently similar to SARS-CoV-2 (96% identity) to cause non-specific alignment [14]. On the other hand, in some cases the virus-specific reads obtained may not be abundant enough for unambiguous detection, which can lead to false negative results. Examples of such cases may be when the viral RNA is highly degraded, or when the sequencing library has been incompletely target enriched by multiple-PCR [15] or hybrid capture [16]. In these scenarios, alignment-based methods may be not specific or sensitive enough. Such alignment-based methods are also computationally intensive, and therefore not particularly fast or efficient.

In order to detect SARS-CoV-2 more quickly and accurately, we have developed an alignment-free method based on k-mer mapping and extension. The use of k-mer-based methods to analyse microbial sequencing data is not a new method. For example, SPINGO [17] provides rapid specie classification for microbial amplicon sequences based on k-mer mapping technology. Kraken2 [18], a very popular taxonomic classification system, is based on k-mer matches. However, we are currently unaware of a fast, reliable, and user-friendly k-mer-based tool for identifying nucleic acids from SARS-CoV-2 and other viruses or microorganisms using sequencing data. This unmet need has led us to develop a new toolset and to provide corresponding k-mer resources.

Here, we present fastv and UniqueKMER, along with the pre-computed unique k-mer resources. The fastv tool has three major functions: to analyse which viral and/or microbial sequences are present in the sequencing data, to determine whether sequences from a specific virus or microorganism (e.g. SARS-CoV-2) can be found in the sequencing data, and to analyse coverage of a specific viral or microbial genome by a set of sequencing data. All three of these methods rely on a unique k-mer set for each microorganism, making it essential to produce high-quality unique k-mer sets. We developed another tool, UniqueKMER, to generate a complete set of unique k-mers for each of a large set of microbial genomes. The unique k-mers can be filtered to remove the k-mers that can be mapped to a reference genome (e.g. the human genome). Generating unique k-mers for tens of thousands of viral and microbial species would typically require tremendous memory and computing resources. We have designed efficient algorithms to make this computation feasible on ordinary computing servers. UniqueKMER requires only one hour to generate reference-filtered unique k-mers for about 12,000 viral genomes on an ordinary PC.

As a lightweight tool, fastv accepts FASTQ data as input, and directly outputs the results in HTML and JSON formats. The HTML result is highly informative and provides interactive reports for manually reading, while the JSON result is structured such that it can easily be used by downstream analysis tools. Prior to the k-mer analysis, fastv automatically performs adapter trimming, quality pruning, base correction, and other pre-processing to ensure the accuracy of k-mer analysis. These pre-processing features are derived from fastp [19], a popular quality control and filtering tool for NGS data previously developed by our group. The fastv tool is ultra-fast - it can process 10M+ bases per second - and can complete the processing of a typical mNGS dataset in a few minutes.

We conducted identification experiments on 27 samples positive for SARS-CoV-2 and 25 samples negative for SARS-CoV-2. The results showed that fastv achieved 100% sensitivity and 100% specificity; and that it can distinguish SARS-CoV-2 from SARS [20], MERS [21], and other coronaviruses [22]. Although our original intention in developing these tools was to quickly identify SARS-CoV-2 from sequencing data, our tool can detect any target virus or microorganism for which a unique k-mer file is provided. We also conducted experiments using several other viral genomes such as Epstein-Barr Virus (EBV), Human Papillomavirus (HPV), and Hepatitis B Virus (HBV). The results demonstrated that our tools perform well on a variety of viral genomic datasets.

## MATERIAL AND METHODS

This section consists of three subsections: rapid identification of microorganisms from sequencing data, algorithms for efficiently generating unique k-mer sets, and pre-generation of unique k-mer sets for common viruses and microorganisms.

### Fastv: rapid identification of microorganisms from sequencing data

Fastv is a highly optimized FASTQ scanner and k-mer mapper. Sequencing reads are first pre-processed to remove adapters and unqualified bases. Continuous k-mers are then computed for each read to be mapped to unique k-mer indexes. Fastv accepts input from any or all of the following three FASTA file types to generate corresponding k-mer indexes:

a. **Unique k-mer collection for a large set of viruses or microorganisms.** This file contains a list of viruses or microorganisms along with their unique k-mers. The identifier of each FASTA entry represents the name of a viral or microbial genome, while its corresponding multi-line sequences represent its unique k-mer keys. This file typically consists of all viruses or microorganisms that might be detected, with reference genomes available.
b. **Unique k-mer set for a specific virus or microorganism**. This file contains a list of k-mer keys unique to a specific virus or microorganism. The sequence of each FASTA entry represents a k-mer key, while its corresponding FASTA identifier represents its position in the viral or microbial genome. This file typically represents the unique k-mer set for the virus or microorganism of interest, such as SARS-CoV-2.
c. **Genome sequences for a specific virus or microorganism.** This file contains one or more reference genomes for the target virus or microorganism. Typically, multiple genomes represent different subtypes of the target virus or microorganism. For instance, if the target virus is HPV, the genomes may comprise HPV-16, HPV-18, HPV-31, etc [23]. Due to the differences among different genome sequences, the coverage and mismatch rate of each genome will be different. This information can be used for microbial subtype identification.

Extraction of k-mers from reads and processing of k-mer indexes are independent processes. Figure 1 summarizes the fastv workflow.

**Figure 1.**
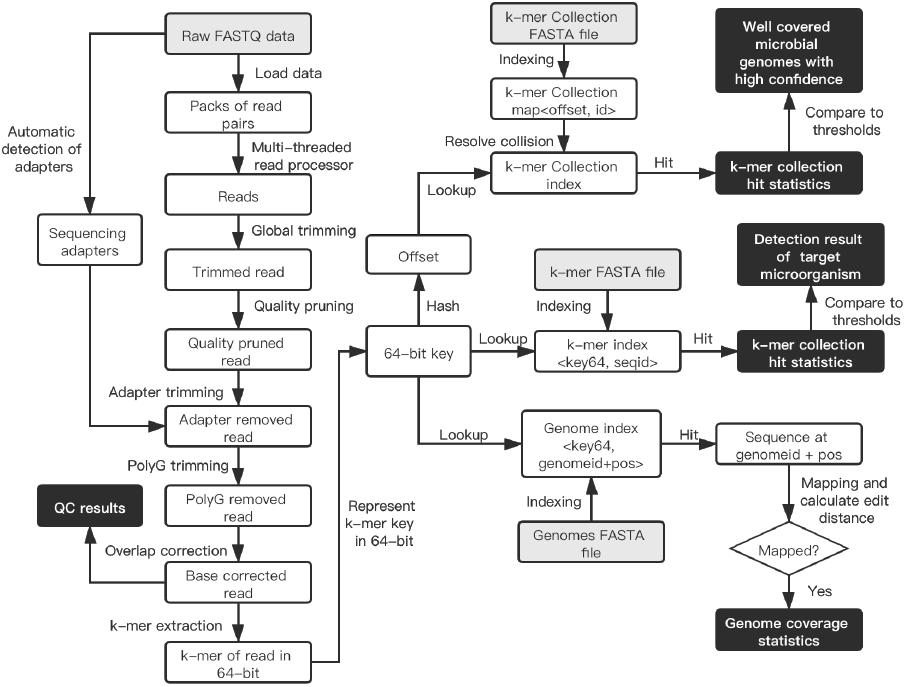
Overview of the fastv workflow. The items with grey backgrounds are input files, while the items with black backgrounds are results that will be output to HTML/JSON reports. An individual thread loads the FASTQ data to read packs (pack size = 1000). Multi-threaded read processors process data pack by pack. For each read or read pair, pre-processing is performed as in fastp. k-mers are extracted from the read and its reverse complement, and are then converted to 64-bit keys. Each 64-bit key is used to search one of the three k-mer indexes built from the k-mer collection file, k-mer file, or genome(s) file.

#### Read pre-processing

To avoid generating erroneous k-mer keys, sequencing adapters and low-quality bases must be removed. Fastv utilizes the adapter cutting and quality pruning features from fastp, which we developed previously. Adapter sequences can either be auto-detected or specified from the command line. For paired-end sequencing data, overlapping regions are detected and incorrect bases in the overlapping regions are corrected. The algorithms for, and implementations of these features can be found in the fastp publication [19]. For data generated by long-read platforms (e.g. PacBio or ONT), long reads are segmented, generating multiple short reads.

#### K-mer generation and representation

To accelerate k-mer generation and lookup, we use a unique 64-bit integer to represent each k-mer. A base (*A*/*T*/*C*/*G*) is represented by two bits, so a 64-bit integer can represent up to a 32-mer, which is sufficiently longfor identifying a virus or microorganism. Any k-mer key that contains a degenerate base (i.e. *N* base) is ignored. This k-mer representation has been used widely in our previous works, such as GeneFuse [24]. A progressive method for k-mer calculation is applied to accelerate k-mer generation. The formula to compute the *n*^*th*^ k-mer key can be denoted as: *kmer*(*n*) = ((*kmer*(*n*-1) << 2) + *base2bit*(*n*)) & *bit*_*mask*, where *base2bit*(*n*) is the 2-bit representation of a base (*A*/*T*/*C*/*G*), and *bit*_*mask* is a 64-bit value determined by the length *k* of k-mer.

#### K-mer collection scanning

The k-mer collection file can contain unique k-mers for tens of thousands of viral or microbial genomes, with each containing hundreds to thousands of k-mer keys. Therefore, there may be tens of millions of k-mer keys in total. It would be trivial to use a map<key, id> to index this data, but such an implementation would result in very slow access. Our approach was to build a hash function to hash the 64-bit key to a larger number (i.e. 2^30^), which stores the index to the k-mer element. Because of the nature of the hash function, two different keys may have the same hash value, resulting in a hash collision [25]. However, as long as the hash function is sufficiently random, the probability of collision will follow a probability distribution that will result in only a small fraction of keys to have hash collisions. This kind of space-for-time approach results in a significant increase in efficiency and makes it feasible to detect tens of thousands of microorganisms at once. The algorithm to build such a k-mer collection index is briefly illustrated as Algorithm 1. It should be noted that, to obtain a higher running speed, we did not use mutex locks to synchronize the k-mer hit counting operation between multiple threads. Our results demonstrate that, while this might lead to slight instability in the results (less than 1 in 10^4^), it does not have an impact on the overall quality of the results.

##### Algorithm 1: generation of KMER collection index

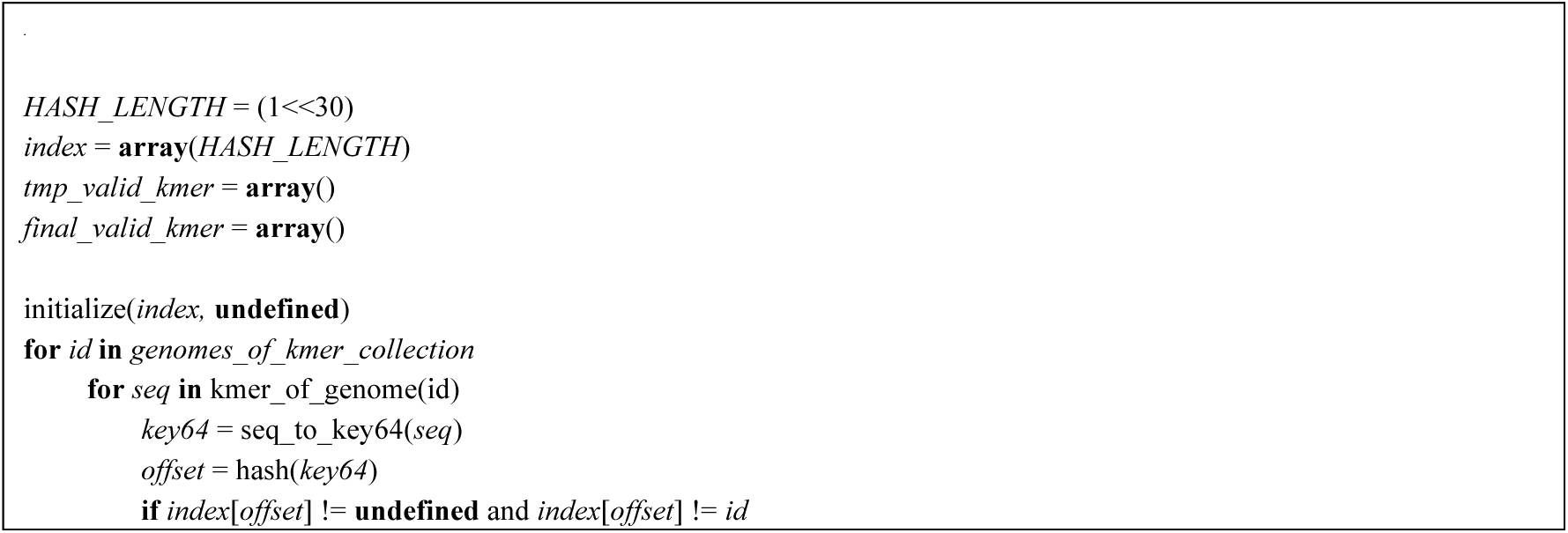
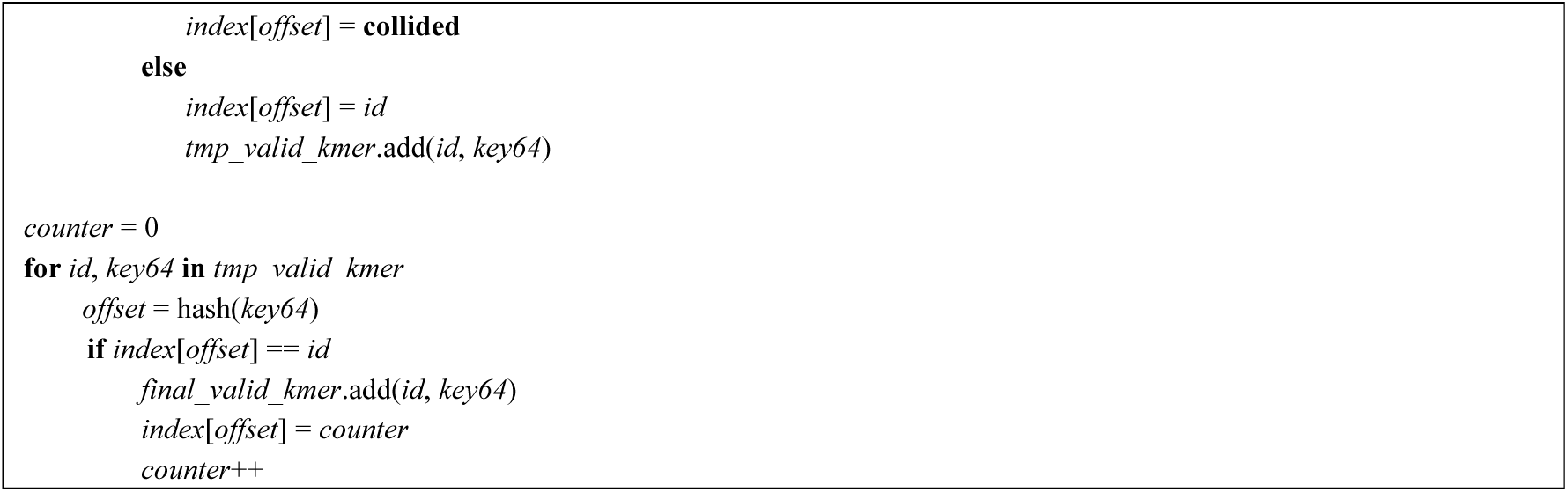

The hash function used in Algorithm 1 is a simple formula that can be calculated efficiently. We have used this hash function in previous works [26]. It utilizes the multiplication and bit manipulation of the key with several big prime numbers:

~~~
**hash**(*key64*) = (1713137323 * ***key64*** + (***key64*** >>12)*7341234131 + (***key64*** >>24)*371371377) & (***HASH_LENGTH***-1)
~~~

#### K-mer scanning for a specific virus or microorganism

Since the k-mer list of a specific virus or microorganism is typically small, it is trivial to implement k-mer scanning on such a short list. A simple unordered map is used to represent the index for such a k-mer set, with the values used as k-mer hit statistics. In contrast to k-mer collection scanning, where multiple threads share a single hit counting statistics array, each thread in the k-mer scanning operation uses its own statistics array. These arrays are subsequently merged to generate the overall statistics, making the k-mer scanning result stable and reproducible.

#### Genome coverage statistics and subtyping

The genomes are indexed as a map, with its key as the same 64-bit k-mer key, and its value as a list of genome positions (GP). A genome position includes a genome ID and a position in that genome. For a given read, if one of its 64-bit k-mer keys is a hit to the genome index, the read will be mapped to the corresponding genome location. The edit distance [27] between the read and the genome sequence will be obtained; and if the edit distance is less than or equal to the threshold, a match will be recorded and the coverage of that genome will be updated. It is worth mentioning that some microbial genomes (i.e. EBV) have a large number of repeated sequences [28], which will result in some keys being hits to many different genome positions. In such cases, fastv divides the coverage and mismatch numbers of a read into multiple parts, and distributes them to each position, producing smooth and uniform coverage. When multiple genomes are input, the coverage result will be sorted by the coverage. This feature can help identify a subtype of a specific virus or microorganism.

#### Visualization

The k-mer scanning results of different inputs are visualized in a figure on a single HTML page. For k-mer scanning of a specific virus or microorganism, the result is simply plotted with the widely used library Plotly.js. For genome k-mer scanning results, we developed a much more efficient toolkit based on native browser utilities to illuminate genome coverage and mismatch ratios. Our highly optimized method can easily support the visualization of hundreds of genomes, which is impossible for most common plotting libraries. A demonstration of a fastv HTML report is shown in Figure 2.

**Figure 2.**
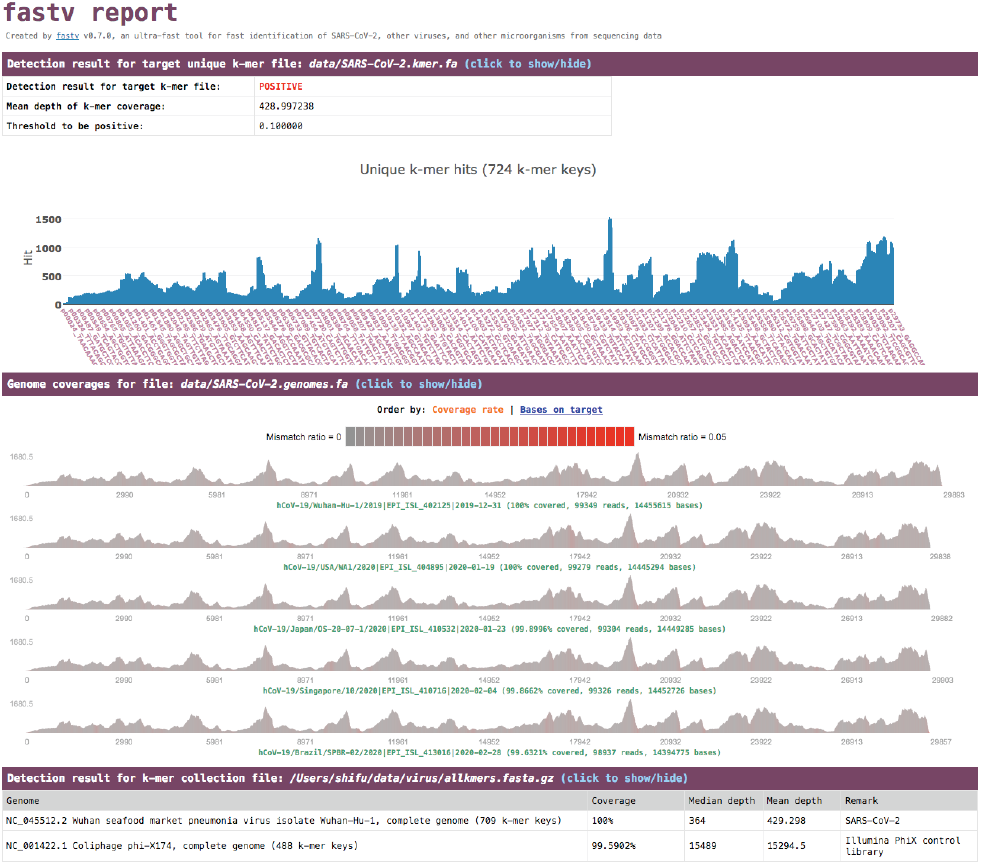
Fastv HTML report demonstration. The result for targeted k-mer hits is visualized using Plotly.js, whereas the result for genome coverage is visualized by a custom toolkit we developed. The k-mer collection scanning result shows that the data contains sequences of two microbial genomes. One is phi-X174, which is actually introduced by the Illumina PhiX control library [29], and the other is SARS-CoV-2. The genome coverage statistics show that SARS-CoV-2 most closely matches strain Wuhan-Hu-1. The red marks indicate the regions with a high mismatch ratio.

### UniqueKMER: efficient unique k-mer generation for large datasets

Since the key features of fastv rely on unique k-mer mapping and extension, it is important to obtain high quality unique k-mer sets for microorganisms of interest. Although a number of k-mer generation tools are currently available [30, 31], none are suitable for our application because we must both generate unique k-mers for tens of thousands of viruses and/or microorganisms, and filter the k-mer keys based on the reference genome. These unmet needs have led us to develop UniqueKMER, a new unique k-mer generation tool. The workflow of UniqueKMER is briefly described in Figure 3.

**Figure 3.**
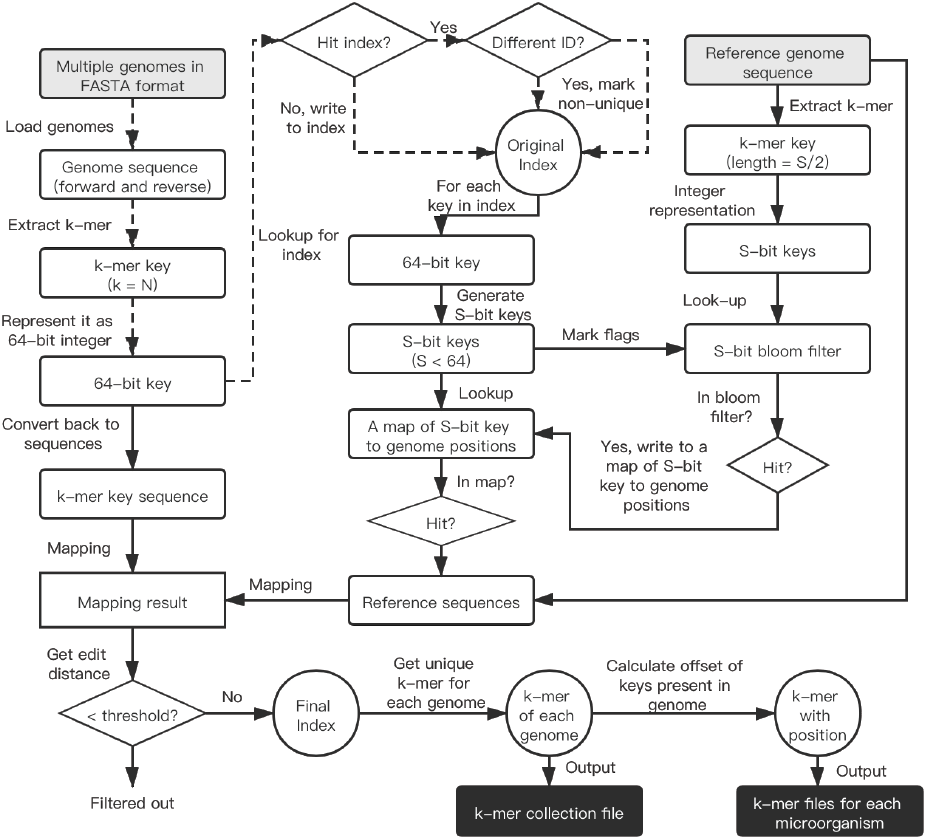
Overview of the UniqueKMER workflow. The items with grey backgrounds are input files, while the items with black backgrounds are output files. The original index consists of unique k-mer keys that belong to a single genome. The original index keys that can be mapped to the reference with edit distance less than the threshold will be filtered out, producing the final index. The S-bit keys are used to identify seeds for key-reference mapping. The k-mer key length *N* is configurable, and is usually set to a number between 20 and 30. The bit length *S* of S-bit is an even number much less than *2N*.

The UniqueKMER workflow consists of two parts (Figure 3): unique k-mer generation and unique k-mer filtering. In the first part all k-mer keys are extracted, and keys that belong to more than one genome are removed as non-unique keys. In the second part, both the keys that exactly match the reference genome and the keys that can be partially mapped to reference genome are removed. Although this can be done with a common aligner such as BWA or Bowtie2, these are not ideal for partial mapping of short sequences. We developed an algorithm based on S-bit seeding and edit distance computation to address this problem, as briefly shown in Algorithm 2.

#### Algorithm 2: unique KMER filtering by reference genome

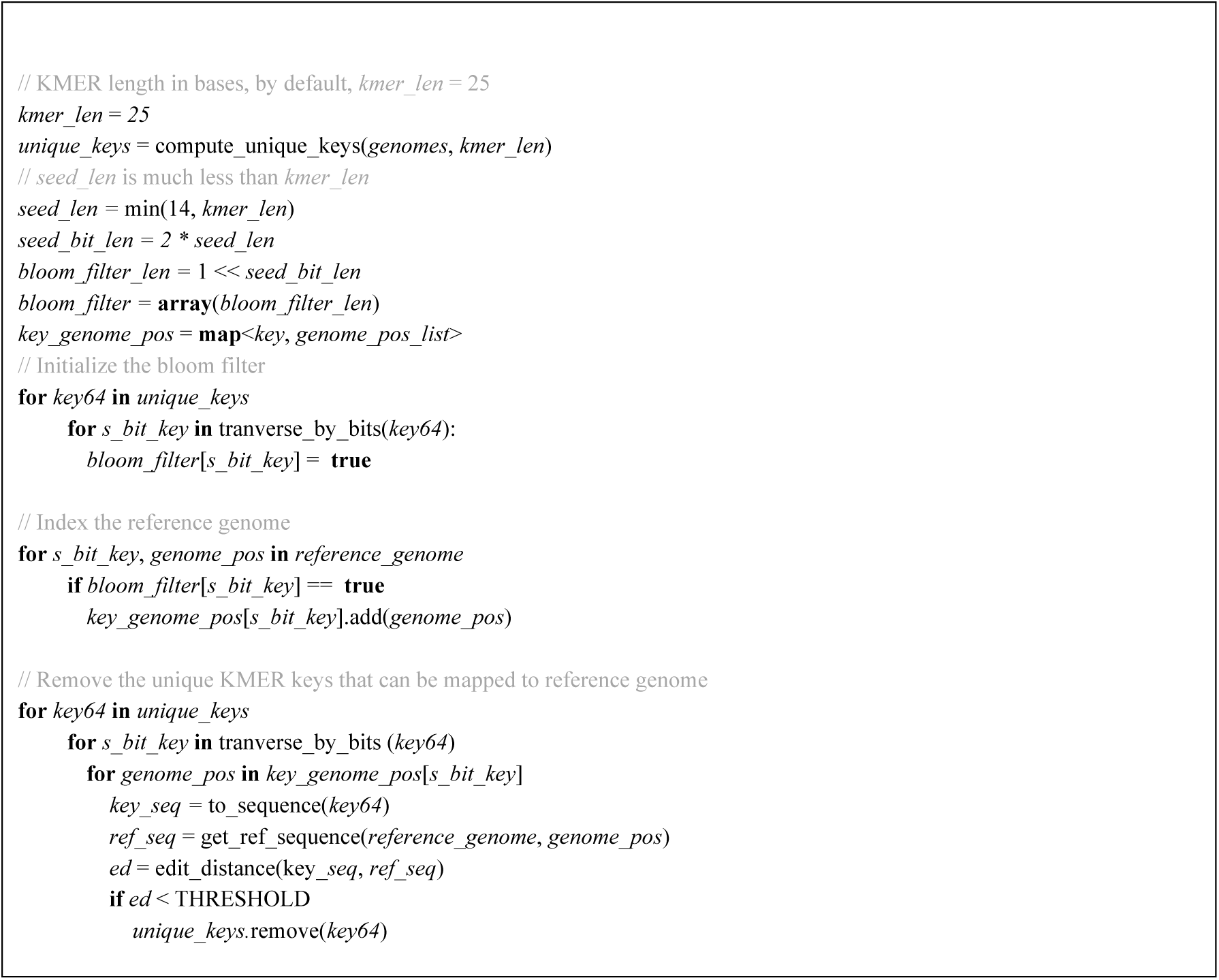

### Pre-generation of unique k-mers for SARS-CoV-2 and other common viruses and microorganisms

We pre-generated unique k-mers for two datasets. The first dataset is the NCBI viral genomes RefSeq database [32], which can be found at https://ftp.ncbi.nlm.nih.gov/refseq/release/viral/. The other pre-generated dataset is the NCBI human bacterial microbiome RefSeq database [33], which can be found at https://ftp.ncbi.nlm.nih.gov/genomes/HUMAN_MICROBIOM/Bacteria/. Because bacterial genomes often have multiple contigs, we concatenated contigs from a single bacterial genome by inserting 32 Ns between each contig, guaranteeing that no artifactual k-mer keys will be introduced unexpectedly. The generated resources can be found at the UniqueKMER repository (https://github.com/OpenGene/UniqueKMER).

SARS-CoV-2 is included in the viral genome list so its unique k-mer set was also generated. We selected twelve SARS-CoV-2 genomes from the GISAID database. Forster et al. recently conducted evolutionary analysis of 160 SARS-CoV-2 genomes [34]. They classified SARS-CoV-2 into three types (A, B, and C) according to amino acid changes, and determined the evolutionary relationships of these types. Based on this work, we selected two ancestral and derived genomes from each type, taking into consideration the date and location of the collected samples. Because SARS-CoV-2 has few mutations to date [35], the similarity between the genomes of different types is very high. Nevertheless, coverage sorting is able to identify the most closely related genome to the query sequence data. The selected SARS-CoV-2 genomes are available at the fastv repository (https://github.com/OpenGene/fastv), and may be updated according to new researches.

## RESULTS

### SARS-CoV-2 identification

To evaluate the performance of fastv for SARS-CoV-2 identification, we conducted experiments on 27 samples that were positive for SARS-CoV-2 and 25 samples that were negative for SARS-CoV-2. The platforms used for sequencing these samples were diverse, and included Illumina, Oxford Nanopore, BGI-Seq, Capillary (Sanger sequencing), and Ion Torrent. For comparison, we also conducted alignment-based identification of SARS-CoV-2 using the widely used aligner BWA. The results are shown in Table 1.

**Table 1.** Comparative performance of fastv and alignment-based method for identification of SARS-CoV-2.

| BioSample Accession # | SRA Accession # | Sequencing Platform | Ground truth of the data | Fastv result | Alignment-based result | Alignment coverage % |
| --- | --- | --- | --- | --- | --- | --- |
| SAMN14422484 | SRP253640 | Ion Torrent | SARS-CoV-2 | POSITIVE | POSITIVE | 56.36 |
| SAMN14381187 | SRP252977 | ONT | SARS-CoV-2 | POSITIVE | POSITIVE | 29.35 |
| SAMN14380341 | SRP252988 | Illumina | SARS-CoV-2 | POSITIVE | POSITIVE | 99.83 |
| SAMN14341640 | SRP251618 | Illumina | SARS-CoV-2 | POSITIVE | POSITIVE | 99.98 |
| SAMN14341639 | SRP251618 | Illumina | SARS-CoV-2 | POSITIVE | POSITIVE | 99.85 |
| SAMN14308026 | SRP251618 | Illumina | SARS-CoV-2 | POSITIVE | POSITIVE | 100.00 |
| SAMN14332760 | SRP252022 | BGISEQ | SARS-CoV-2 | POSITIVE | POSITIVE | 36.20 |
| SAMN14308029 | SRP251618 | Illumina | SARS-CoV-2 | POSITIVE | POSITIVE | 99.41 |
| SAMN14180202 | SRP250653 | Illumina | SARS-CoV-2 | POSITIVE | POSITIVE | 100.00 |
| SAMN14154205 | SRP250294 | Illumina | SARS-CoV-2 | POSITIVE | POSITIVE | 99.91 |
| SAMN14154204 | SRP250294 | ONT | SARS-CoV-2 | POSITIVE | POSITIVE | 100.00 |
| SAMN14154203 | SRP250294 | Illumina | SARS-CoV-2 | POSITIVE | POSITIVE | 48.24 |
| SAMN14154202 | SRP250294 | ONT | SARS-CoV-2 | POSITIVE | POSITIVE | 100.00 |
| SAMN14154201 | SRP250294 | Illumina | SARS-CoV-2 | POSITIVE | POSITIVE | 44.49 |
| SAMN14154200 | SRP250294 | ONT | SARS-CoV-2 | POSITIVE | POSITIVE | 100.00 |
| SAMN14154199 | SRP250294 | Illumina | SARS-CoV-2 | POSITIVE | POSITIVE | 99.88 |
| SAMN14154198 | SRP250294 | ONT | SARS-CoV-2 | POSITIVE | POSITIVE | 100.00 |
| SAMN14082199 | SRP249613 | Illumina | SARS-CoV-2 | POSITIVE | POSITIVE | 97.60 |
| SAMN14082197 | SRP249613 | Illumina | SARS-CoV-2 | POSITIVE | POSITIVE | 99.96 |
| SAMN14082196 | SRP249613 | Illumina | SARS-CoV-2 | POSITIVE | POSITIVE | 92.57 |
| SAMN14082200 | SRP249613 | Illumina | SARS-CoV-2 | POSITIVE | POSITIVE | 99.91 |
| SAMN13922059 | SRP245409 | Illumina | SARS-CoV-2 | POSITIVE | POSITIVE | 99.44 |
| SAMN13872787 | SRP242226 | Illumina | SARS-CoV-2 | POSITIVE | POSITIVE | 100.00 |
| SAMN13872786 | SRP242226 | Illumina | SARS-CoV-2 | POSITIVE | POSITIVE | 100.00 |
| SAMN13871323 | SRP242169 | ONT | SARS-CoV-2 | POSITIVE | POSITIVE | 99.78 |
| SAMN13898864 | SRP242169 | ONT | SARS-CoV-2 | POSITIVE | POSITIVE | 100.00 |
| SAMN14445407 | SRP253783 | BGISEQ | SARS-CoV-2 | POSITIVE | NEGATIVE | 13.97 |
| SAMN14086238 | SRP249478 | Illumina | Bat Coronaviruses | NEGATIVE | NEGATIVE | 0.00 |
| SAMN14086235 | SRP249478 | Illumina | Bat Coronaviruses | NEGATIVE | NEGATIVE | 0.50 |
| SAMN14086234 | SRP249478 | Illumina | Bat Coronaviruses | NEGATIVE | NEGATIVE | 0.00 |
| SAMN14086233 | SRP249478 | Illumina | Bat Coronaviruses | NEGATIVE | NEGATIVE | 0.00 |
| SAMN14086230 | SRP249478 | Illumina | Bat Coronaviruses | NEGATIVE | NEGATIVE | 0.00 |
| SAMN14082201 | SRP249482 | Illumina | Bat coronavirus RaTG13 | NEGATIVE | POSITIVE | 96.04 |
| SAMN02688745 | SRP040070 | Illumina | MERS-CoV | NEGATIVE | NEGATIVE | 0.00 |
| SAMN02688792 | SRP040070 | Illumina | MERS-CoV | NEGATIVE | NEGATIVE | 0.00 |
| SAMN02688873 | SRP040070 | Illumina | MERS-CoV | NEGATIVE | NEGATIVE | 0.12 |
| SAMN02402894 | SRP033021 | Illumina | SARS-CoV | NEGATIVE | NEGATIVE | 17.71 |
| SAMN02402954 | SRP033022 | Illumina | SARS-CoV | NEGATIVE | NEGATIVE | 17.50 |
| SAMN02402960 | SRP033023 | Illumina | SARS-CoV | NEGATIVE | NEGATIVE | 5.57 |
| SAMEA5841282 | ERP116617 | ONT | Human coronavirus 229E | NEGATIVE | NEGATIVE | 0.00 |
| SAMEA5841281 | ERP116617 | ONT | Human coronavirus 229E | NEGATIVE | NEGATIVE | 0.00 |
| SAMEA5841278 | ERP116617 | ONT | Human coronavirus 229E | NEGATIVE | NEGATIVE | 0.13 |
| SAMEA3931206 | ERP014917 | Illumina | Human coronavirus OC43 | NEGATIVE | NEGATIVE | 0.00 |
| SAMEA3931205 | ERP014917 | Illumina | Human coronavirus NL63 | NEGATIVE | NEGATIVE | 0.00 |
| SAMEA3931204 | ERP014917 | Illumina | Human coronavirus HKU1 | NEGATIVE | NEGATIVE | 0.00 |
| SAMEA3931214 | ERP014917 | Illumina | Human respirovirus 1 | NEGATIVE | NEGATIVE | 0.00 |
| SAMEA3931208 | ERP014917 | Illumina | Human bocavirus 1 | NEGATIVE | NEGATIVE | 0.00 |
| SAMEA3931225 | ERP014917 | Illumina | Respiratory syncytial virus | NEGATIVE | NEGATIVE | 0.00 |
| SAMN03386977 | SRP056027 | Illumina | HKU5 Coronavirus | NEGATIVE | NEGATIVE | 12.44 |
| SAMN03386974 | SRP056027 | Illumina | Bat coronavirus HKU3 | NEGATIVE | NEGATIVE | 0.63 |
| SAMN12113789 | SRP204847 | Capillary | Influenza A Virus HKU11 | NEGATIVE | NEGATIVE | 0.00 |
| SAMN12113790 | SRP204848 | Capillary | Influenza A Virus HKU12 | NEGATIVE | NEGATIVE | 0.00 |

As shown in Table 1, fastv achieved 100% sensitivity and 100% specificity for all tested samples; and could distinguish SARS-CoV-2 from SARS, MERS, and other coronaviruses. The pipeline for alignment-based SARS-CoV-2 identification, which is described in Supplementary File 1, is based on the widely used alignment method BWA-MEM [12]. The alignment-based pipeline failed to identify a SARS-CoV-2 sample, which was previously enriched by using multiplex PCR technology. It also incorrectly identified the bat coronavirus RaTG13 sample as SARS-CoV-2 since genome of RaTG13 has about 96% similarity to genome of SARS-CoV-2 [14]. Even if careful manual adjustment of the alignment parameters may result in a better result on this dataset, it is difficult to ensure that the adjusted parameters can achieve good results on other datasets. The fastv results are based on the default parameters determined before this experiment. Therefore results obtained using fastv are more robust and reliable than those obtained using the BWA alignment-based method.

### Identification of other viruses

We also conducted experiments using data from other viruses and microorganisms. Epstein-Barr virus (EBV) has a long repetitive region within its genome, which often causes difficulties for other k-mer-based algorithms because these algorithms map a k-mer key to the location where it first appears. But our optimized k-mer mapping algorithm distributes hits corresponding to a k-mer key to all places where it appears, resulting in smoother coverage. An example of EBV identification is shown in Figure 4 to illustrate the smooth coverage obtained for genomes with repetitive regions.

**Figure 4.**
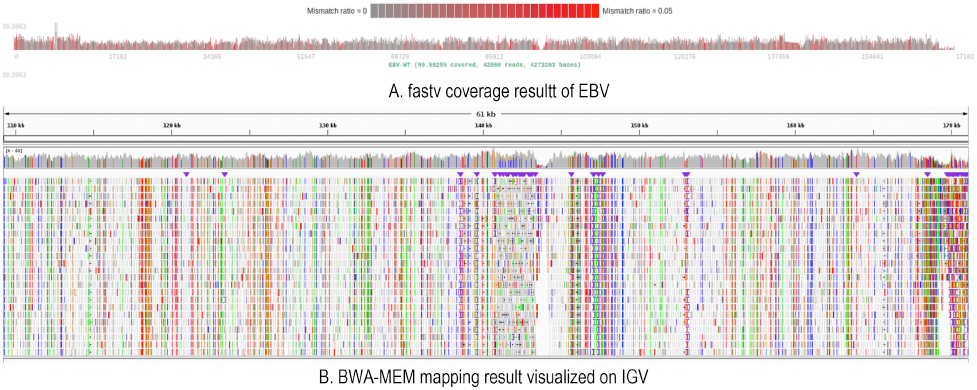
EBV identification using fastv. The EBV genome has a large repetitive region between 12kb and 35kb. The coverage result generated by fastv (A) shows even coverage of this region, and is similar to the IGV visualization of the BWA alignment result (B). The data used in this experiment was whole genome sequencing of an EBV-positive sample, downloaded from NCBI SRA (SRA accession: ERR1293949).

We evaluated the subtyping function of fastv with sequencing data from samples infected with HPV and HBV. For HPV, genomes of nine HPV subtypes (HPV-6b, HPV-11, HPV-16, HPV-18, HPV-31, HPV-33, HPV-45, HPV-52, HPV-58) were included in the genomes FASTA file, and three datasets (SRA accessions: SRR1609138,SRR160913 and SRR1609140) were tested. For HBV, genomes of eight HBV subtypes (HBV-A to HBV-H) were included and four datasets (SRA accessions: SRR11308108, SRR11308109, SRR11308111 and SRR11308112) were tested. The results showed that fastv could distinguish the virus subtypes for all these data very accurately. Figure 5 shows the correct HBV subtyping result of dataset SRR11308112, which is the output of sequencing data from a HBV-C infected sample.

**Figure 5.**
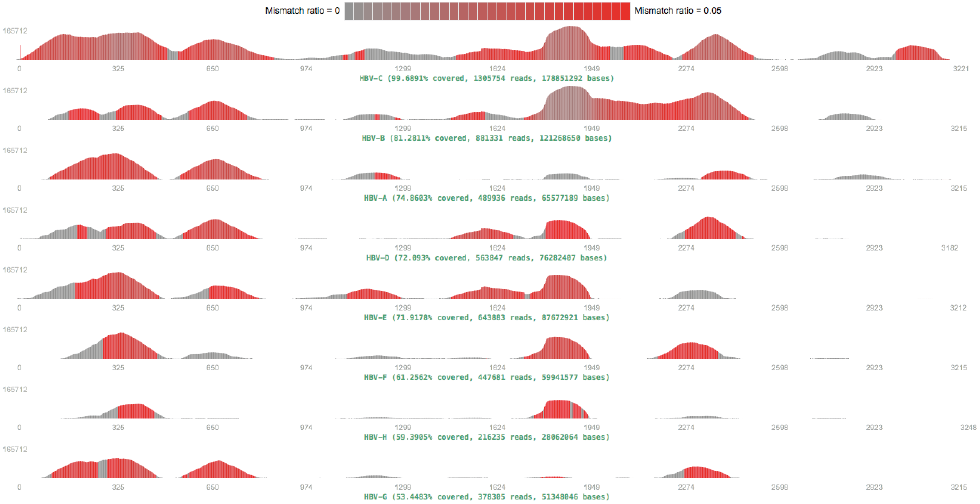
HBV subtyping using fastv. Eight HBV subtypes (HBV-A to HBV-H) are included in the genome list. The subtyping result is HBV-C, which is 99.69% covered. HBV genome has many conservative regions [36], where the genomes of different subtypes are very similar. This led to the other HBV subtypes appearing to be partly covered, but obviously not covered as well as HBV-C.

### Identification of pathogen from mNGS data without set a target virus or microorganism

When the pathogen is unknown, fastv can also be used to quickly identify microorganisms in sequenced samples to speculate what the pathogen is. In order to use this function, a k-mer collection file containing many possible viruses and microorganisms should be first prepared. For the convenience of users, we have pre-generated a k-mer collection file, which consists of genomes for all viruses and human bacteria that with a reference genome provided in NCBI RefSeq database. This pre-generated k-mer collection file can be downloaded from fastv repository. After scanning the FASTQ data, fastv will report the k-mer coverage for each microbial genome with valid hits. This information can be used to identify the possible pathogen. We evaluated seven mNGS datasets (SRA accessions: SRP006887, SRP006881, SRP000376, SRP007321, ERS4389819, SRP000657 and SRP004485), which were generated by Illumina and LS454 sequencers. All the correct pathogens were successfully detected, with the k-mer coverage varying from 15.19% to 99.94%.

## DISCUSSION

In summary, we describe a new tool, fastv, for rapid identification of viruses and microorganisms from sequencing data. This tool is based on the k-mer mapping and extension method, and relies on high-quality unique k-mers. We also describe a new tool, UniqueKMER, to generate such high-quality unique k-mer sets for a large collection of viruses and microorganisms. Experimental results show that with the k-mers generated by UniqueKMER, fastv is able to detect SARS-CoV-2 with 100% sensitivity and 100% specificity.

Because of the rapid and unpredictable spread of COVID-19, it is important to develop inexpensive, rapid, and reliable methods for identification of its causative agent, SARS-CoV-2. Next generation sequencing-based methods are suitable for SARS-CoV-2 detection, and offer some advantages over other detection methods. Therefore, computational tools that can rapidly and reliably identify SARS-CoV-2 from sequencing data will be valuable to the research community. Fastv can also output on-target (e.g. SARS-CoV-2) clean reads to individual FASTQ files, which can be input to downstream analysis pipelines. For example, genome assembly with the on-target clean reads will be simpler and faster. The on-target reads can also be input to database search utilities like BLAST [37].

Although our original intention in developing fastv and UniqueKMER was to quickly identify SARS-CoV-2 from sequencing data, these tools can be used more generally to detect any target virus or microorganism for which unique k-mer files are provided. The results of our experiments with EBV, HPV, and HBV sequencing data demonstrate the general applicability of our tools. Because fastv can rapidly scan tens of thousands of genomes, it is a powerful tool for analysing mNGS data, particularly for identifying pathogens from mNGS data. Fastv can be used for rapid screening of possible viruses or bacteria to get the information that can further help determine the pathogen. We will continue to update our resource library so that researchers can directly use the pre-generated high-quality unique k-mer files for mNGS data analysis. It should be pointed out that currently fastv sorts the results according to the genome coverage and median k-mer hits, without considering the pathogenicity of each virus or microorganism. In future work, we will further link the microbial pathogen databases, like the FDA-ARGOS [38], to provide better functions for pathogen identification.

## Availability

As part of the OpenGene projects, fastv and UniqueKMER are open-sourced through the MIT license. Fastv is available at https://github.com/OpenGene/fastv, and UniqueKMER is available at https://github.com/OpenGene/UniqueKMER. The pre-computed unique k-mer resources are also provided in these repositories.

## Key Points

This tool presents a new tool fastv for rapid identification of SARS-Cov-2, other viruses and microorganisms. Another tool UniqueKMER is presented for generation of high-quality unique k-mers. Unique k-mer resources for tens of thousands of viruses and microorganisms have been pre-computed, and uploaded to the tools’ repositories.

## Supplementary Data

A pipeline for alignment-based SARS-CoV-2 identification was provided in **Supplementary file 1**.

## Acknowledgement

The authors would like to thank the community users for testing these tools and reporting bugs.

~~~
**# Use fastp for adapter trimming and quality pruning**
**fastp** -i $sample_R1.fastq.gz -o $sample_clean_R1.fastq.gz -I $sample_R2.fastq.gz -O
$sample_clean_R2.fastq.gz -h $sample_clean.html -j $sample_clean.json > $sample_clean.log &
**# Extract microbial reads by filtering human reads.**
**bwa** mem -R “@RG\tID:$sample\tLB:$sample\tPL:$sample\tSM:$sample” -t 8 -k 32 -M
$path/bwaindex/hg19.fa $sample_clean_R1.fastq.gz $sample_clean_R2.fastq.gz | **sentieon** util sort -o $sample.hg19.sort.bam -t 8 --sam2bam -i -
**samtools** view -b -f 12 -F 256 $sample.hg19.sort.bam > $sample.hg19.bothEndsUnmapped.bam
**samtools** sort -@ 8 -n $sample.hg19.bothEndsUnmapped.bam -o
$sample.hg19.bothEndsUnmapped.sorted.bam
**bamToFastq** -i $sample.hg19.bothEndsUnmapped.sorted.bam -fq $sample.rmhost.clean.1.fq -fq2
$sample.rmhost.clean.2.fq
**# Remove Ribosomal RNA (rRNA) reads**
**bwa** mem -R “@RG\tID:$sample\tLB:$sample\tPL:$sample\tSM:$sample” -t 8 -k 32 -M
$path/rRNA_fa/sequence.fa $sample.rmhost.clean.1.fq $sample.rmhost.clean.2.fq | **sentieon** util sort -o $sample.rRNA.sort.bam -t 8 --sam2bam -i -
**samtools** view -b -f 12 -F 256 $sample.rRNA.sort.bam > $sample.rRNA.bothEndsUnmapped.bam
**samtools** sort -@ 8 -n $sample.rRNA.bothEndsUnmapped.bam -o
$sample.rRNA.bothEndsUnmapped.sorted.bam
**bamToFastq** -i $sample.rRNA.bothEndsUnmapped.sorted.bam -fq $sample.rmrRNA.clean.1.fq
-fq2 $sample.rmrRNA.clean.2.fq
**# Align reads to the SARS-CoV-2 genome, remove duplicates, and calculated the coverage.**
**bwa** mem -R “@RG\tID:$sample\tLB:$sample\tPL:$sample\tSM:$sample” -t 8 -k 32 -M
$path/bwaindex/SARS-CoV-2.fa $sample.rmrRNA.clean.1.fq $sample.rmrRNA.clean.2.fq |
**sentieon** util sort -o $sample.sort.bam -t 8 --sam2bam -i -
**gencore** -i $sample.sort.bam -o $sample.gencore.sort.bam -r $path/bwaindex/SARS-CoV-2.fa -j
$sample.gencore.json -h $sample.gencore.html
**samtools** index $sample.gencore.sort.bams
**samtools** depth -aa $sample.gencore.sort.bam > $sample.gencore.sort.depth
**samtools** mpileup -AB -Q 25 -q 30 -d 100000 -f $path/bwaindex/SARS-CoV-2.fa
$sample.gencore.sort.bam > $sample.gencore.sort.mpileup
# **Determine the result by evaluate the coverage of SARS-CoV-2 genome, greater than 20% means positive.**
#!/bin/bash
genome_length=$(awk ‘{print NR}’ $sample.gencore.sort.depth | tail -n1)
coverd_base=$(awk ‘{print NR}’ $sample.gencore.sort.mpileup | tail -n1)
percent=$(awk ‘BEGIN{printf “%0.2f”,(‘$coverd_base’/’$genome_length’)*100}’)
awk -v num1=$percent -v num2=20 ‘BEGIN{print(num1>num2)?”POSITIVE”:”NEGATIVE”}’
~~~

